# Naturalized *Dolichogenidea gelechiidivoris* Marsh (Hymenoptera: Braconidae) complement the resident parasitoid complex of *Tuta absoluta* (Meyrick) (Lepidopera:Gelechiidae) in Spain

**DOI:** 10.1101/2021.05.27.445932

**Authors:** Carmen Denis, Jordi Riudavets, Oscar Alomar, Nuria Agustí, Helena Gonzalez-Valero, Martina Cubí, Montserrat Matas, David Rodríguez, Kees van Achterberg, Judit Arnó

## Abstract

Our study aimed to assess the contribution of natural parasitism due to *Necremnus tutae* Ribes & Bernardo (Hymenoptera: Eulophidae) to the biological control of *Tuta absoluta* (Meyrick) (Lepidopera:Gelechiidae) in commercial plots where an IPM program based on the use of predatory mirid bugs was implemented. During the samplings, the presence of another parasitoid was detected and, therefore, a second part of our study intended to identify this species and to evaluate the importance of its natural populations in the biological control of the pest. Leaflets with *T. absoluta* galleries were collected during 2017–2020 from commercial tomato plots in the horticultural production area of Catalonia (Northeast Spain), including greenhouses, open fields, and roof covered tunnels that lack side walls. In the laboratory, *T. absoluta* larvae were classified as ectoparasitized, alive, or dead. Reared parasitoids from ectoparasitized larvae were mostly morphologically identified as *Necremnus* sp. with parasitism rates that peaked in summer months with values between 9 and 15%. Some of these ectoparasitized larvae also yielded another parasitoid identified as *Dolichogenidea gelechiidivoris* Marsh (Hymenoptera: Braconidae) by both morphological and molecular-DNA barcoding methods. In 2020, parasitism rates due to *D. gelechiidivoris* that increased with season up to 22%. Our work reports for the first time in Europe the presence of the neotropical species *D. gelechiidivoris* adding this biocontrol agent to the resident parasitoid complex of *T. absoluta* in Spain.

## Introduction

*Tuta absoluta* (Meyrick) (Lepidopera:Gelechiidae) is native to South America where has been considered an important tomato pest since a long time (Larrain 1987). The first report outside its area of origin was from Spain in 2006 (Urbaneja et al. 2007). From then onwards, the pest did spread over the Mediterranean basin, and then quickly colonized Africa and Asia (Desneux et al. 2010; Desneux et al. 2011), threatening tomato production (Biondi et al. 2018).

Although insecticides still remain the main control tool in many world areas, many efforts have targeted sustainable biological control methods (Biondi et al. 2018). In Spain, the positive role of the predatory mirid bugs *Macrolophus pygmaeus* (Rambur) and *Nesidiocoris tenuis* (Reuter) (Hemiptera: Miridae) was soon acknowledged (Arnó et al. 2009; Urbaneja et al. 2009). Successful IPM programs based on both predators already in use at the time of invasion did greatly contribute to the control of *T. absoluta* (Urbaneja et al. 2012). These two bugs remain as cornerstones for the biological control of several pests in the area (Arnó et al. 2018).

Surveys of parasitoids that could complement the poor predator action on *T. absoluta* larvae were soon undertaken in the Mediterranean (Zappalà et al. 2013; Gabarra et al. 2014), and several larval parasitoids of *T. absoluta* within the Eulophidae, Braconidae, Chalcididae, Ichneumonidae and Pteromalidae were recorded (Biondi et al. 2018; Mansour et al. 2018). Out of these species, *Necremnus tutae* Ribes & Bernardo (Hymenoptera: Eulophidae), first identified as *Necremnus* nr. *artynes*, was found consistently parasitizing *T. absoluta* (Gebiola et al. 2015), and several studies have recognized the contribution of natural populations to the biological control of the pest (Abbes et al. 2014; Crisol-Martínez and van der Blom 2019; Arnó et al. in press). This parasitoid attracted the attention of the biocontrol industry, and for some time it was commercially available, but the high host-killing rate was a serious drawback for successful mass rearing (Calvo at al. 2016) and production has been discontinued.

To further assess the contribution of *Necremnus tutae* to the biological control of *T. absoluta*, our first goal was to estimate the natural parasitism in commercial plots where the IPM program based on the use of predatory mirid bugs was implemented. However, since we observed the presence of another parasitoid, the second part of our study aimed to identify this species and to evaluate the importance of its natural populations in the biological control of *T. absoluta*.

## Materials and Methods

Samples were collected during 2017–2020 from commercial tomato plots in the horticultural production area of Catalonia (Northeast Spain), including greenhouses, open fields, and roof covered tunnels that lack side walls (Table 1). Each plot was sampled up to 21 times depending on crop duration and *T. absoluta* infestation levels. Plots were sampled by randomly walking in the plot and collecting leaflets that had galleries big enough to host a second to third instar larva of *T. absoluta*. Sampling terminated after 20 minutes or a maximum of 25 leaflets, whichever was reached first, in order to minimize sampling time and costs, particularly when infestation levels were low (Naranjo 2008).

**Table 1:**
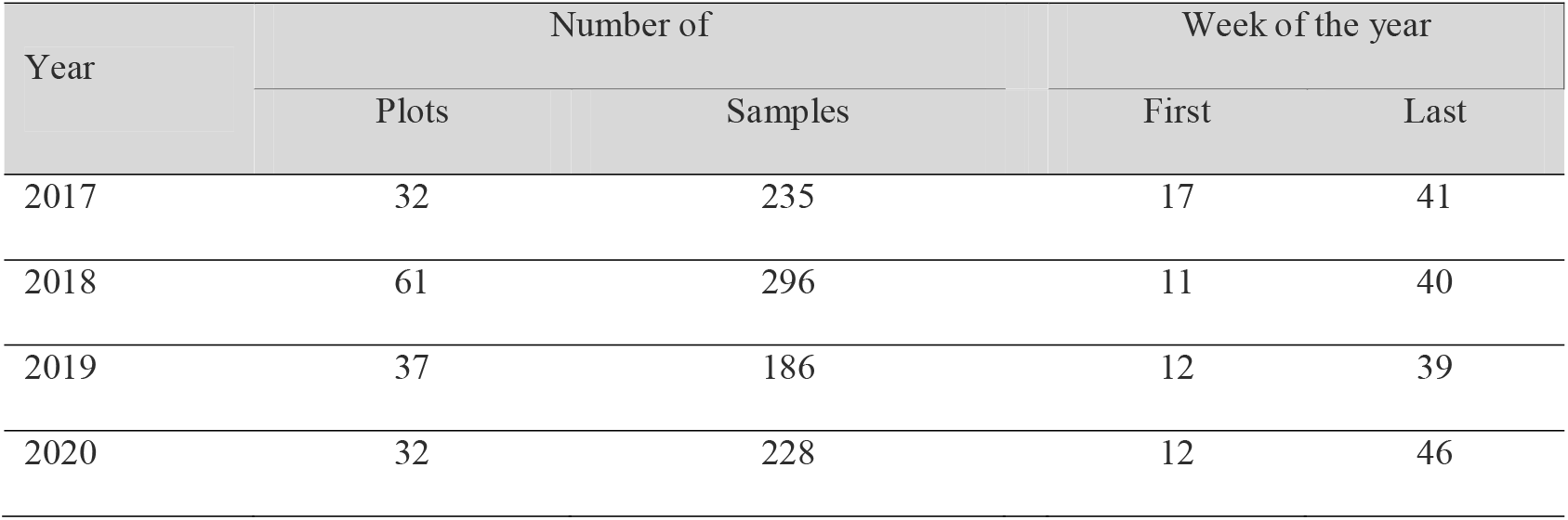
Number of plots surveyed, number of samples that had leaflets with at least one *T. absoluta* larva, and week of the year when first and last samples were taken. Each sample corresponds to a maximum of 25 leaflets during up to 20 minutes from one crop on one date.

Leaflets were taken to the laboratory and inspected under a stereomicroscope. *Tuta absoluta* larvae were classified as “ectoparasitized” (with pupae, larvae or eggs of a parasitoid on the *T. absoluta* larva), “alive” (not ectoparasitized and able to crawl when touched with a fine brush), or “dead” (not ectoparasitized and unable to crawl when touched with a fine brush), and the number of each category was recorded. As our initial interest was on the contribution of *N*.*tutae* to the control of *T. absoluta*, during 2017, 2018 and 2019, only the ectoparasitized larvae were placed inside Petri dishes (maximum of 3 larvae per dish), and kept at room temperature for at least 2-3 weeks.

During 2019, the recurrent presence in the samples, albeit at a low rate, of a parasitoid belonging to the Braconidae was observed. Therefore, additional samples were collected from nine tomato fields in September. From these samples, alive, dead and ectoparasitized larvae were individualized in Petri dishes and kept in the climatic chamber (25°C) for a maximum of 42 days until the emergence of *T. absoluta* or of adult parasitoids.

From the samples collected in 2020, the number of alive, dead, and ectoparasitized *T. absoluta* larvae was also recorded as before, but only larvae that were alive, with no clearly visible ectoparasitoids, were kept in order to determine the parasitization rate by endoparasitoids. Larvae were placed in a Petri dish (maximum of 3 larvae per dish) together with the leaflet and stored at room temperature for a maximum of 3 weeks until the pupation of *T. absoluta* or the emergence of a parasitoid. Emerged parasitoids from the four years samples were collected and stored in 70% alcohol for further identification.

Due to the high irregularity of infestation levels in the plots, the number of leaflets collected from each plot and date (a sample) was very variable, and furthermore, not all the galleries had a *T. absoluta* larvae inside. Consequently, the total number of larvae (alive, dead or ectoparasitized) collected in each plot was very variable, and sometimes very low. Leaflets with only empty galleries and no *T. absoluta* larvae were discarded. To summarize the levels of parasitism, samples were pooled for each month and year. The percentage of monthly ectoparasitism was calculated by dividing the number of ectoparasitized larvae by the total number of larvae recorded each month. Additionally, for the 2020 samples of alive *T. absoluta* larvae, monthly parasitism due to endoparasitoids was calculated dividing the number of emerged parasitoids by the total number of larvae recorded each month.

Adult parasitoids were first identified to family and sub-family level using the keys by Grissell and Schauff (1990) and Hanson and Gauld (2006). Eulophidae were further identified to genus using the keys by Askew (1968) and Gebiola et al. (2015). Microgastrinae (Hymenoptera: Braconidae) were first identified to genus following the descriptions of Fernandez-Triana et al. (2020), and then as *Dolichogenidea gelechiidivoris* Marsh (=*Apanteles gelechiidivoris*) using the description of Marsh (1975). Additional material examined for morphological identification were 10 ♂♂ and 10 ♀♀, from a laboratory rearing started with adults that emerged from *T. absoluta* larvae collected in 2019 from several tomato fields in El Maresme county (31TDF49 to 31TDG92, Catalonia, Spain). These voucher specimens were prepared and deposited in Naturalis Biodiversity Center (Leiden, The Netherlands).

To confirm the morphological identification of *D. gelechiidivoris*, 11 specimens were sequenced for DNA barcoding identification. The specimens emerged from samples collected in different locations from eastern Catalonia along a transect of 100 Km (from 31TDF17 to 31TDG84 and 31TEG03, in the municipalities of Viladecans (2018), Mataró (2019, 2020), Argentona (2018), Santa Susana (2017, 2019, 2020), Blanes (2017), Fornells de la Selva (2019), and Calonge (2018). An additional specimen from a previous study (Arnó et al. in press) collected in 2016 in Malgrat de Mar was also DNA barcoded.

Total genomic DNA was extracted from whole insects by using SpeedTools Tissue DNA Extraction Kit (Biotools, Madrid, Spain) following the manufacturer protocol and eluted in 100 μl of BBE buffer provided by the manufacturer and stored at -20°C. A 658-bp region of the CO1 gene was amplified with the following protocol using primers LepF1 5-ATTCAACCAATCATAAAGATATTGG-3 and LepR1 5-TAAACTTCTGGATGTCCAAAAAATCA-3 (Smith *et al*. 2006). PCR reaction volumes (20 µl) contained 2 µl of resuspended DNA, 10 µl of Master Mix (Biotools, Madrid, Spain) and 0.4 µl of each primer [10 µM]. Samples were amplified in a 2720 thermal cycler (Applied Biosystems, CA, USA) using the thermocycling profile of one cycle of 2 min at 94°C; five cycles of 40 sec at 94°C, 40 sec at 45°C, and 1 min at 72°C; followed by 35 cycles of 40 sec at 94°C, 40 sec at 51°C, and 1 min at 72°C, with a final step of 5 min at 72°C (Smith *et al*. 2006). PCR products were analysed by electrophoresis in 2.4% agarose gels stained with GelRed® (Biotium, Hayward, CA) and visualized under UV light. They were purified with QIAquick PCR Purification kit (Qiagen) and bidirectionally sequenced by using BIGDYE 3.1 on an ABI 3730 DNA Analyzer (Applied Biosystems) at the Genomics Unit of the Barcelona Science Park (University of Barcelona). Obtained sequences were compared against the reference database *Barcode of Life Data System* (BOLD, http://www.boldsystems.org/), to find the matching species.

## Results

DNA barcoding successfully confirmed the initial identification based on morphological characters of the 11 analyzed specimens as *D. gelechiidivoris*, regardless of location and year of collection. Obtained similarity percentages ranged from 100 to 99,48% compared with the 13 *D. gelechiidivoris* available sequences at the time of the analysis (February 2021) in the GenBank database (Accession codes: KX443088, HQ558975-HQ558977, JN282071-JN282078 and JQ849955).. Obtained sequences were also deposited to the GenBank database (Accession codes: MZ298974-MZ298984).

Monthly levels of ectoparasitized larvae over the four sampling years are displayed in Table 2. They were recorded from April to November, with levels ranging from 0.1% (May 2017) to 35.7% (November 2020). Apart from this exceptionally high value, each year the parasitism peaked during the summer months of August and September with values between 9 and 15%.

**Table 2.** Percentages of ectoparasitized *T. absoluta* larvae recorded from tomato leaflets collected in commercial tomato plots. Percentages were calculated as the number of ectoparasitoized larvae over the total number of larvae collected per month. In brackets are the number of samples that had leaflets with at least one *T. absoluta* larvae.

| Month | % ectoparasitized larvae |  |  |  | *T. absoluta* larvae (num. samples) |  |  |  |
| --- | --- | --- | --- | --- | --- | --- | --- | --- |
|  | 2017 | 2018 | 2019 | 2020 | 2017 | 2018 | 2019 | 2020 |
| Mar | - | 0 | 0 | 0 | - | 480 (25) | 113 (12) | 11 (3) |
| Apr | 0 | 0.8 | 1.0 | 0 | 32 (5) | 612 (47) | 308 (22) | 14 (6) |
| May | 0.1 | 3.0 | 5.0 | 0 | 714 (50) | 833 (52) | 360 (25) | 148 (19) |
| Jun | 0.6 | 7.3 | 6.2 | 0.3 | 519 (52) | 575 (43) | 389 (27) | 398 (48) |
| Jul | 4.9 | 5.2 | 4.3 | 0.6 | 409 (53) | 515 (55) | 531 (45) | 530 (43) |
| Aug | 10.6 | 5.5 | 13.5 | 1.8 | 256 (31) | 455 (48) | 319 (36) | 610 (47) |
| Sep | 11.6 | 14.6 | 7.6 | 9.2 | 242 (27) | 213 (21) | 197 (19) | 370 (40) |
| Oct | 9.1 | 3.1 | - | 7.4 | 186 (16) | 65 (5) | - | 202 (17) |
| Nov | - | - | - | 35.7 | - | - | - | 14 (5) |

Most parasitoids that emerged from ectoparasitized larvae from 2017 to 2019 were Eulophids (87%) and a smaller percentage (11%) belonged to Braconidae (Table 3). Among the Eulophidae, eight genera were recorded: *Necremnus* (162 individuals), *Pnigalio* (7), *Neochrysocaris* (5), *Diglyphus* (4), *Stenomesius* (4), *Aprostocetus* (3), *Cirrospilus* (2), and *Sympiesis* (1). Out of the 25 individuals belonging to Braconidae that emerged from larvae that had an ectoparasitoid on them (Table 3), 23 were morphologically identified as *D. gelechiidivoris* same as the four individuals collected in 2016.

**Table 3:** Number of adult parasitoids of different Hymenoptera families reared from ectoparasitized *T. absoluta* larvae in samples collected from commercial plots in the different years of sampling.

| Hymenoptera family | Year of sampling |  |  |
| --- | --- | --- | --- |
|  | 2017 | 2018 | 2019 |
| Eulophidae | 23 | 68 | 98 |
| Braconidae | 5 | 8 | 12 |
| Torymidae | 1 | 0 | 0 |
| Platygastridae | 1 | 0 | 0 |
| Diapriidae | 0 | 1 | 0 |
| Aphelinidae | 0 | 0 | 1 |

The additional samples collected in September 2019 yielded 170 *T. absoluta* larvae. From 21 ectoparasitized *T. absoluta* larvae emerged 11 *Necremnus* sp., 2 *Diglyphus* sp., 1 *Neochrysocharis* sp., and 1 unidentified Eulophid. From 114 dead *T. absoluta* larvae emerged 6 *Necremnus* sp., and from the 35 alive larvae emerged 13 *D. gelechiidivoris*.

In 2020, 1,872 alive larvae were collected from 228 samples from 32 plots. From these larvae, we obtained a total of 264 parasitoids that emerged from 92 different samples collected from 20 plots. A subsample of 165 were morphologically identified as *D. gelechiidivoris*, together with one *Neochrysocharis* sp., and one from the subfamily Alysiinae. As can be observed in Figure 1, the percentage of endoparasitism steadily increased from May until October. No endoparasitoids were recorded in March, April and November.

**Figure 1:**
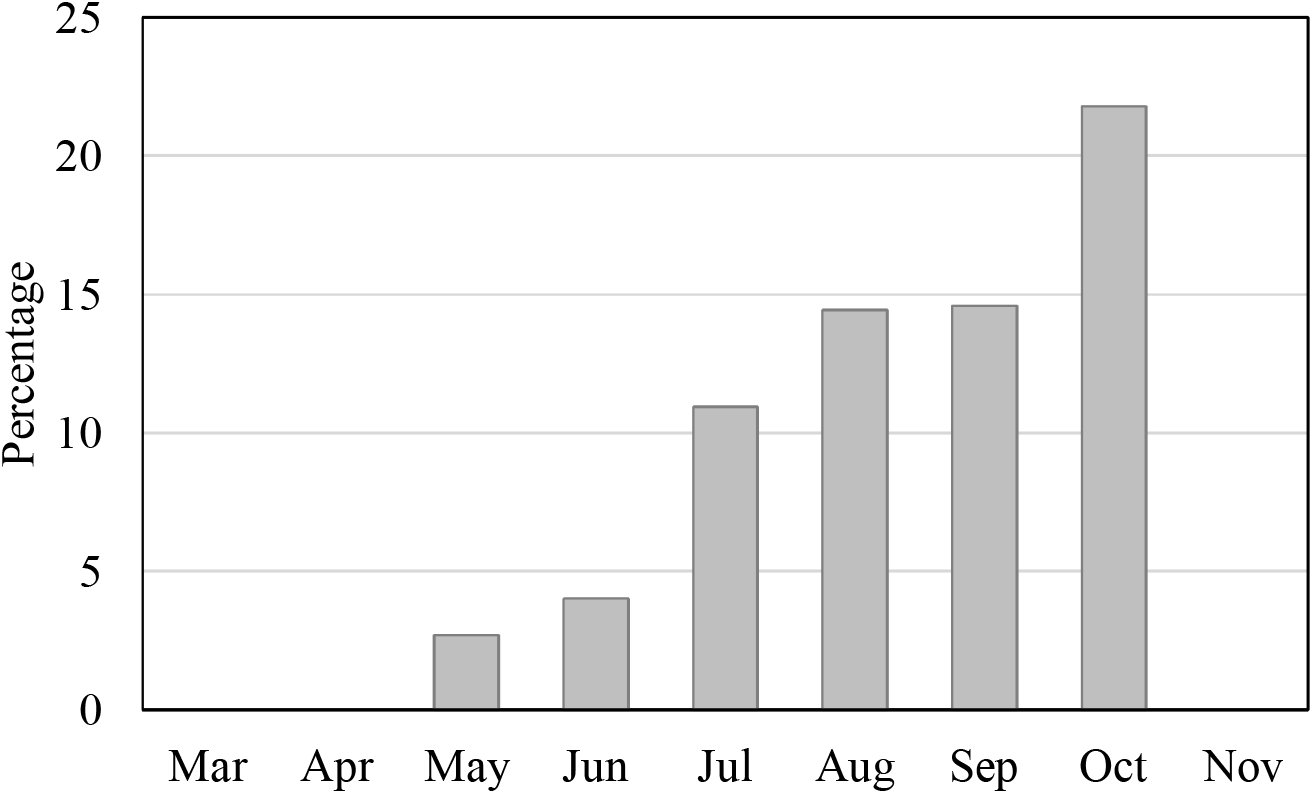
Monthly percentage of endoparasitized *T. absoluta* larvae in commercial tomato plots in 2020. Percentages were calculated as the number of emerged parasitoids over the total number of larvae recorded per month. Total number of *T. absoluta* larvae and number of samples are displayed in Table 2.

## Discussion

Results of our four-years field survey confirmed the relevance of Eulophidae as larval parasitoids of *T. absoluta* in the Mediterranean basin. All genera except *Aprostocetus* had been previously reported (Biondi et al. 2018). We reared three individuals of this genus from a single sample collected in September 2017. *Aprostocetus* are parasitic or hyperparasitic in Lepidoptera (Sakaltaş and Tüzün 2014), and Mirchev et al. (2001) refer to an *Aprostocetus* sp. as an hyperparasitoid of a Braconid parasitizing a Gelechiid.

Among Eulophidae, our results corroborate the importance of *Necremnus* sp. as the most widespread ectoparasitoid of *T. absoluta* larvae in the Mediterranean, as reported before (Ferracini et al. 2012; Zappalà et al. 2013; Gabarra et al. 2014; Gebiola et al. 2015; Biondi et al. 2018). In fact, *N. tutae*, which is by far the predominant *Necremnus* species in the area of our survey (Arnó et al. in press), was considered a promising parasitoid to be released for *T. absoluta* control (Calvo et al. 2013; Chailleux et al. 2014; Bodino et al. 2016; Calvo et al. 2016), although currently it is not commercially available. Rates of ectoparasitized larvae found in the present study, mostly due to *Necremnus* sp., were similar to those recorded in other field studies in the Mediterranean basin. In the same area of the present study, parasitism rates by *N. tutae* in sentinel plants were close to 20% (Arnó et al. in press), and in Tunisia it was of 26% in sentinel plants, and between 11 and 15% when sampling the crop (Abbes et al. 2014). Much higher rates, up to 73%, were recorded in tomato greenhouses in the southeast of Spain (Crisol-Martínez and van der Blom 2019).

Contribution of *N. tutae* to the control of *T. absoluta* goes further than only parasitization. As many Eulophidae, it kills more larvae than parasitizes in order to obtain nutrients that have a strong positive effect on its reproduction (Ferracini et al. 2012; Balzan and Wäckers 2013; Calvo et al. 2013; Chailleux et al. 2014; Calvo et al. 2016; Bodino et al. 2019). In the present study, although the number of dead larvae was recorded, we could not determine the exact causes of mortality because even all sampled plots were managed according to an IPM program based on predatory mirids, insecticide applications were also occasionally required.

An important outcome of our study was the detection of *D. gelechiidivoris* in field collected samples from 2017 to 2020, and also in one sample collected in 2016. *Dolichogenidea gelechiidivoris* is native of South America, where it is considered as an important agent of biological control (Larrain 1987, Agudelo and Kaimowitz 1997; Vallejo 1999; Salas-Gervasio et al. 2019). In Colombia, mass rearing protocols were developed to release this parasitoid for *T. absoluta* control (Morales et al. 2013). Furthermore, in 2017 it was imported to Kenia from Peru to contribute to the control of *T. absoluta* in Africa (Aigbedion-Atalor et al. 2020). Another species, *Dolichogenidea appellator* (Telenga) (= *Dolichogenidea litae* (Nixon)), was occasionally found parasitizing *T. absoluta* in the same area as the present study (Gabarra et al., 2014), and was found also associated with this pest in Sudan (Mansour et al. 2018).

To our knowledge, this is the first report of *D. gelechiidivoris* naturally occurring outside its area of origin. Since there is no record of intentional introduction of *D. gelechiidivoris* into Europe, and the importation to Africa took place in 2017, one year after our first detection of this species in 2016, the results of our survey suggest that this parasitoid was unintentionally introduced from the Neotropics together with the pest. Accidental introductions of natural enemies in new territories are not strange. Roy et al (2011) stated that most of the alien arthropod predator and parasite species in Europe arrived accidentally, as part of worldwide movement of invasive pests that is facilitated by global trade. Trade of infested fruits has been pointed out as the most probable cause of the arrival of *T. absoluta* to Spain from South America and the quick spread of the pest (Desneux et al. 2010).

As all Microgastrinae, *D. gelechiidivoris* is a koinobiont solitary larval endoparasitoid (Fernandez-Triana et al. 2020), and the host remains alive until the end of the parasitoid development. Of the 2017–2019 samples of *T. absoluta* larvae, we only kept those that had an ectoparasitoid egg or larva on them, but 11.5% of the emerged parasitoids were *D. gelechiidivoris* (Table 3). This suggests that there is no clear recognition of previous parasitism between ectoparasitoids (mainly *Necremnus* sp.), and *D. gelechiidivoris*.

When we maintained all alive larvae to record parasitoid emergence (additional 2019 samples, and all 2020 samples), the endoparasitism rate was 7.6% in September 2019 and increased up to 22% from May to October 2020 (Fig. 1). However, the real rate of parasitism by *D. gelechiidivoris* in 2020 was probably underestimated since about 11% of ectoparasitized larvae collected between 2017 and 2019 yielded *D. gelechiidivoris*, and ectoparasitized larvae of 2020 had been discarded.

In this scenario of coexistence of several natural enemies, the interactions between *D. gelechiidivoris* and *Necremnus sp*. will be of great importance. For example, young females of the ectoparasitoid *Dineulophus phthorimaeae* De Santis (Hymenoptera: Eulophidae) avoided multiparasitism on the microgastrinid endoparasitoid *Pseudapanteles dignus* (Muesebeck) (Hymenoptera: Braconidae), but older females did not discriminate heterospecific parasitized *T. absoluta* larvae, and joint action of both parasitoids exerted an important control of *T. absoluta* (up to 80% of host larvae mortality) (Salas-Gervasio et al., 2019). The outcome of competition between parasitoids attacking the same host depends on many factors that may explain the dominance of one parasitoid over another, e.g. where the venom of idiobiont ectoparasitoids has little or no effect on the development of endoparasitic koinobionts (Harvey 2013), although in multiple parasitisms between an ectoparasitoid and an endoparasitoid, the former normally wins (Mitsunaga and Yano 2004). Furthermore, the interaction between *D. gelechiidivoris* and predatory mirids will be also of interest. These predators prefer to prey on eggs but may also feed on young *T. absoluta* larvae (Arnó et al. 2009; Urbaneja et al. 2009), which are the preferred host instar of the parasitoid (Aigbedion-Atalor et al. 2020). *Nesidiocoris tenuis* did not prey on nor did reduce the oviposition by *D. gelechiidivoris*, and the efficacy of both natural enemies together on *T. absoluta* was significantly higher than either natural enemy alone (Aigbedion-Atalor 2020). The outcome of the interactions among these biocontrol agents will be determinant for a more successful control of *T. absoluta* (Tarusikirwa et al. 2020).

## Acknowledgements

The present research was supported by the Spanish Ministry of Economy and Competitiveness (AGL2016-77373-C2-1-R) and the Ministry of Agriculture, Livestock, Fisheries and Food of the Generalitat de Catalunya. Authors from IRTA were supported by the CERCA Programme/Generalitat de Catalunya. Carmen Denis was supported by a PhD grant of BECAL-PY. We are in debt to Dr. Valmir Antonio Costa (Instituto Biológico of Campinas, Brazil) and Dr. José Fernandez-Triana (Canadian National Collection of Insects, Ottawa, Canada) for their advice in the identification of parasitoids.

